# Stress-induced neuronal hypertrophy decreases the intrinsic excitability in stress habituation

**DOI:** 10.1101/593665

**Authors:** Sara Matovic, Aoi Ichiyama, Hiroyuki Igarashi, Eric W Salter, Xie-Fan Wang, Mathilde Henry, Nathalie Vernoux, Marie-Eve Tremblay, Wataru Inoue

**Author notes:** **Corresponding Author:** Wataru Inoue.

## Abstract

A rapid activation of the hypothalamic-pituitary-adrenal (HPA) axis is a hallmark stress response to an imminent threat, but its chronic activation can be detrimental. Thus, the long-term survival of animals requires experience-dependent fine-tuning of the stress response. However, the cellular mechanisms underlying the ability to decrease the stress responsiveness of the HPA axis remain largely unsolved. Using a stress habituation model in male mice and slice patch-clamp electrophysiology, we studied hypothalamic corticotropin-releasing hormone neurons that form the apex of the HPA axis. We found that the intrinsic excitability of these neurons substantially decreased after daily repeated restraint stress in a time course that coincided with their loss of stress responsiveness *in vivo*. This plasticity of intrinsic excitability co-developed with an expansion of surface membrane area, resulting in an increase in input conductance with minimal changes in conductance density. Moreover, multi-photon and electron microcopy data found that repeated stress augmented ruffling of the plasma membrane, suggesting an ultrastructural plasticity that efficiently accommodates membrane area expansion with proportionally less expansion of gross cell volume. Overall, we report a novel structure-function relationship for intrinsic excitability plasticity that correlates with habituation of the neuroendocrine stress response.

**Significance statement:** The long-term survival of animals requires experience-dependent fine-tuning of stress response. Using a mouse model of repeated stress that develops habituation of the hypothalamic-pituitary-adrenal (HPA) axis, our study demonstrates a robust decrease in the intrinsic excitability of the output neuroendocrine neurons of the HPA axis. Mechanistically, we show that repeated stress increases the cell size of these neurons (i.e. surface membrane area). This cell-size change increases input conductance, and hence decreases excitability. Our findings challenge a conventional view that plasticity of intrinsic excitability relies on changes on membrane excitability resulting from up- and down-regulation of various voltage-gated ion channels. Our study reports a novel structure-function relationship for intrinsic excitability plasticity that correlates with habituation of the neuroendocrine stress response.

## Introduction

Activation of the hypothalamic-pituitary-adrenal (HPA) axis is a hallmark of the stress response conserved across vertebrates, culminating in widespread actions of glucocorticoids (Denver, 2009). Although adaptive in the short-term, protracted recruitment of this energetically costly response can be maladaptive (Munck et al., 1984; Sapolsky et al., 2000; McEwen, 2007). Indeed, the HPA axis is exquisitely flexible and fine-tunes its stress sensitivity based on the learned significance of the stressor. For example, in both humans and experimental animals, the HPA axis normally responds less and less (i.e. habituates) to repeated encounters of psychological stress (Aguilera, 1994; Chen and Herbert, 1995; Kirschbaum et al., 1995; Bhatnagar et al., 2002; Kim and Han, 2006; Uchida et al., 2008). By contrast, the lack of habituation is associated with stress vulnerability and contributes to maladaptive consequences of chronic stress (McEwen and Seeman, 1999; Epel et al., 2000; Kudielka et al., 2006; Uchida et al., 2008; Gianferante et al., 2014). Despite the biological and clinical importance of HPA axis habituation, surprisingly little is known about the neural plasticity mechanisms through which repetition of stressful experiences refines the sensitivity of the HPA axis to the stressor.

The output of the HPA axis depends on the release of corticotropin releasing hormone (CRH) from neuroendocrine neurons in the paraventricular nucleus of the hypothalamus (PVN). Previous studies show that the hormonal habituation to repeated restraint, a primarily psychological stressor, is paralleled by a reduced up-regulation of immediate early gene c-Fos in PVN-CRH neurons upon stress re-exposure (Ma et al., 1999; Stamp and Herbert, 1999; Cole et al., 2000; Carter et al., 2004; Girotti et al., 2006). These data indicate that plasticity mechanisms, which control the stress sensitivity of PVN-CRH neurons, are important for the HPA axis habituation. While this may be partly achieved by changes in brain areas upstream of the PVN (Bhatnagar et al., 2002; Girotti et al., 2006), emerging evidence also shows that the PVN itself is highly labile and undergoes various types of plasticity after both acute and chronic stress (Bains et al., 2015; Herman and Tasker, 2016). Thus, the PVN offers an entry point to identify and examine the cellular and molecular mechanisms of plasticity that may contribute to HPA axis habituation. Considering that all stress-relevant signals for the neuroendocrine outputs eventually converge onto and translate into the activity of PVN-CRH neurons (Ulrich-Lai and Herman, 2009), plasticity at the level of these neurons would have an important control over the stress sensitivity of the HPA axis.

Here, we report that the intrinsic excitability of PVN-CRH neurons is strongly attenuated over 21 days of daily repeated restraint, in a time course that coincides with their loss of stress responsiveness *in vivo* in mice. Our data collectively indicates that the decrease in PVN-CRH neurons’ intrinsic excitability is due to an increase in surface membrane area (i.e. cell size), which consequently increases the input (whole-cell) conductance with little changes in the conductance density (conductance per unit membrane area). Our findings identify a new form of structure-function relationship involved in intrinsic excitability plasticity as well as a neuronal correlate for HPA axis habituation.

## Materials and Methods

### Animals

All experimental procedures were approved by the University of Western Ontario Animal Use Subcommittee and University Council on Animal Care in accordance with the Canadian Council on Animal Care guidelines. Homozygous *crh-IRES-Cre* (Stock No: 012704, the Jackson Laboratory) and *Ai14* (Stock No: 007908, the Jackson Laboratory) mice were mated, and the resulting heterozygous crh-IRES-Cre;Ai14 offspring were used as CRH-reporter mice as characterized in detail previously (Wamsteeker Cusulin et al., 2013; Itoi et al., 2014; Chen et al., 2015). The bright tdTomato expression allows for visual identification of CRH neurons in brain slices prepared for immunohistochemistry and *ex vivo* electrophysiology recordings. All animals used were male and between 8-12 weeks old at the time of sacrifice. They were group housed (2-4 mice per cage) in a standard shoebox mouse cage supplied with a mouse housing, paper nesting materials and wood chip bedding. The mice were housed on a 12/12 hour light/dark cycle (lights on 07:00) in a temperature-controlled (23 ± 1°C) room with free access to food and water. Cages were cleaned every 7 days.

### Stress protocols

For restraint stress, mice were placed in a restrainer constructed from a 50 mL conical tube with multiple ventilation holes. In order to restrict the forward/backward movement of mice, the inner space length of the restrainer was adjusted by moving a disc wall that forms the head end of the restrainer. For repeated restraint stress, the one hour restraint stress session was repeated for 7 or 21 consecutive days (between 13:00-14:00). For stress recovery, a group of mice were first subjected to 21 days of repeated restraint stress and then left undisturbed in their home cage for 7 days. The control mice were kept in their home cage in the animal housing facility.

### Immunohistochemistry

C-Fos expression was examined immediately after a 1 hour-long episode of restraint stress that was given for the first time (Acute), one day after 7 or 21 (1 hour-long) episodes of daily repeated restraint (7RRS or 21RRS), or one day after 7 days of a no stress recovery period following the 21RRS (Recovery). As a control, we also studied stress-naïve (Control) mice (**Figure 1A**). The mice were deeply anaesthetized with sodium pentobarbital (100 mg/kg, intraperitoneally). A tail-pinch test was performed to ensure the depth of the anesthesia. The mice were then transcardially perfused with ice-cold saline solution (0.9% NaCl) followed by 4% paraformaldehyde (PFA, Sigma) dissolved in phosphate-buffered saline (PBS, pH 7.4). The brains were removed and fixed in 4% PFA at 4°C overnight. The brains were then coronally sectioned into 50 μm thick slices using a vibratome (VT1000S, Leica Biosystems, Concord, ON, Canada). Sections were stored in a cryoprotectant solution (30% glycerol, 30% ethylenglycol, in 20 mM PB) at ‒20°C until use. Immunohistochemistry was performed on free-floating sections. Sections were rinsed in PBS 3 times for 5 minutes each and then incubated in a blocking solution (3% normal donkey serum, 0.3% Triton X-100 and 0.03% NaN_3_ in PBS) for 1 hour. Slices were then incubated with anti-c-Fos rabbit monoclonal antibody (Cell Signalling Technology, cat: 2250S, 1:1000 dilution in blocking solution) for overnight at room temperature. After three washes in PBS, the sections were incubated with Alexa Fluor 647 donkey anti-rabbit IgG (ThermoFisher Scientific, cat: A-31573, 1:500 dilution in blocking solution). After three washes with PBS, the sections were incubated in 4’,6-diamidino-2-phenylindole (DAPI, 100 ng/ml in PBS, Sigma, cat: D9542) for 10 minutes. After two washes with PBS, the sections were mounted on glass slides and cover slipped using Fluoromount-G mounting medium (Electron Microscopy Sciences (EMS), cat: 17984-25).

**Figure 1.**
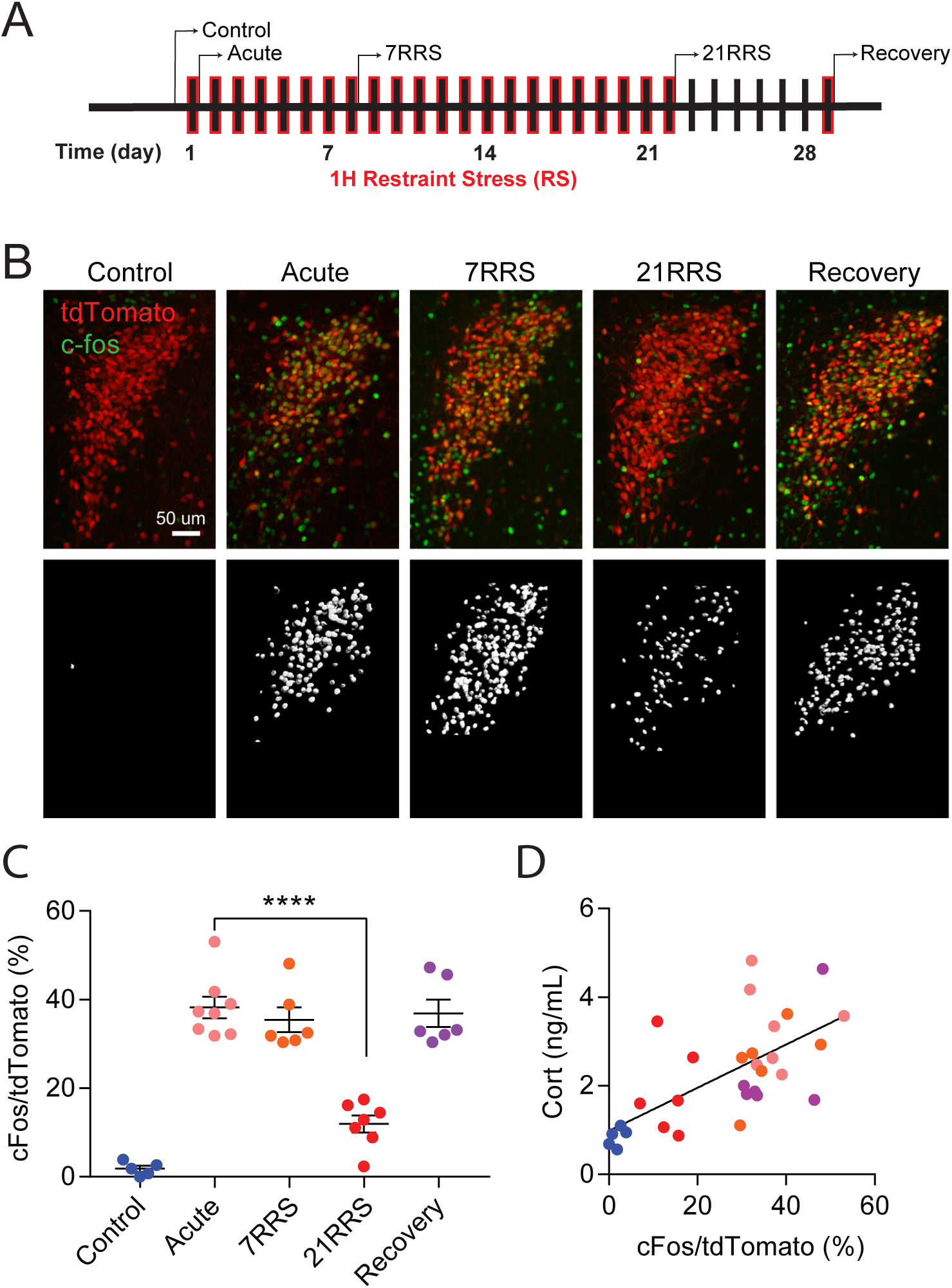
Restraint stress-induced c-Fos expression in PVN-CRH neurons diminishes after 21RRS, but not 7RRS. **(A)** Experimental timeline. Mice received daily, 1-hour-long restraint stress (red) and were perfused immediately after the stress. Control mice were perfused without restraint stress. **(B)** Top: examples of c-Fos immunohistochemistry (green) and tdTomato expression (red) in the PVN of CRH reporter mice. Bottom: co-localization (white) of c-Fos and tdTomato. **(C)** Summary graph for c-Fos/tdTomato volume (%). Acute restraint strongly increased c-Fos expression in tdTomato-positive PVN-CRH neurons as compared to control. 21RRS, but not 7RRS, diminished the restraint-induced c-Fos expression. The loss of c-Fos induction spontaneously recovered after a 7 day no-stress recovery period. **** p < 0.0001. **(D)** C-Fos expression in PVN-CRH neurons significantly correlated with circulating corticosterone levels. Error bars are S.E.M.

C-Fos immunohistochemistry sections were imaged and analyzed as follows. Z stack images (0.685 µm thick optical sections, 20-23 sections) were obtained with a confocal microscope (Leica SP8, Leica-Microsystems) using a 20x objective (HC PL APO CS2, 0.75 NA, dry, Leica-Microsystems). Experimenters were blinded for treatments prior to image analysis. Each image was reconstructed in 3D and quantified using Imaris (Imaris v7.6.4, Bitplane, AG, Zurich, Switzerland). To estimate the volume positive for tdTomato and c-Fos expression, automatic thresholding was applied and a surface was created for each channel. The same threshold values were then used to create a colocalization channel dual-positive for tdTomato and c-Fos. A third surface was then created for the colocalization channel, determining the volume of the colocalization. The degree of cFos expression by tdTomato-positive CRH neurons was derived from colocalization volume divided by tdTomato volume for individual images.

### Reconstruction of biocytin-filled neurons

To estimate the size of electrophysiologically recorded cells, biocytin (0.5 % w/v, ThermoFisher Scientific) was included in the internal solution. The slices were fixed in 4% PFA for 24 hours at 4°C. The filled cells were then visualized by incubating slices with streptavidin-Alexa 488 (1:500, ThermoFisher Scientific) diluted in PBS containing 2% TritonX-100. Dye-filled cells were imaged with a confocal microscope (Leica SP8, Leica-Microsystems) using a 63x (HC PL APO CS2, 1.4 NA oil-immersion, Leica-Microsystems) objective lens with optical section thickness at 0.3 μm. The confocal Z-stack images were 3D reconstructed using Imaris software and surface membrane area was calculated by applying automatic thresholds. Experimenters were blinded for treatments prior to image analysis. To measure the cell soma size, a cubic region of interest was applied to encompass only the soma. The diameter of proximal dendrite was measured at 2 µm from the perceived end of the cell soma by using the measurement point function in Imaris software.

### Electrophysiology

For the electrophysiology experiments, mice were sacrificed on the next day after the last restraint stress (e.g. for 21RRS, on day 22 without restraint stress on the day of sacrifice). Control mice were sacrificed without any stress, and acute stress mice were restraint stressed and sacrificed immediately after the end of stress. The mice were deeply anesthetized with isoflurane and decapitated. Brains were then quickly removed from the skull and placed in icy slicing solution containing (in mM): 87 NaCl, 2.5 KCl, 25 NaHCO_3_, 0.5 CaCl_2_, 7 MgCl_2_, 1.25 NaH_2_PO_4_, 25 glucose and 75 sucrose (Osmolarity: 315-320 mOsm), saturated with 95% O_2_/5% CO_2_. 250 µm thick coronal sections containing the PVN were cut using a vibratome (VT1200 S, Leica). Sections were then placed in artificial cerebral spinal fluid (aCSF) containing (in mM): 126 NaCl, 2.5 KCl, 1.25 NaH2PO_4_, 26 NaHCO_3_, 10 glucose, 2.5 CaCl_2_ and 1.5 MgCl_2_ (Osmolarity: 295-300 mOsm), saturated in 95% O_2_/5% CO_2,_ maintained at 36°C for 30 minutes, and thereafter kept at room temperature in the same aCSF for the rest of the day.

PVN slices were transferred to a recording chamber superfused with aCSF (flow rate 1-2 mL/min, 28-32°C). Slices were visualized using an upright microscope (BX51WI, Olympus) equipped with infrared differential interference contrast and epifluorescence optics as well as a digital camera (Rolera-XR, Q-Imaging). CRH neurons were identified by their expression of TdTomato. Patch clamp recording pipettes were pulled from borosilicate glass (Cat#: BF150-86-10, Sutter Instrument) using a Flaming/Brown Micropipette Puller (P-1000, Sutter Instrument) to a tip resistance of 3-5 MΩ. Electrodes were filled with an internal solution containing (in mM): 108 K-gluconate, 2 MgCl_2_, 8 Na-gluconate, 8 KCl, 1 K_2_-ethylene glycol-bis(β-aminoethyl ether)-N,N,N’,N’-tetraacetic acid (EGTA), 4 K_2_-ATP, 0.3 Na_3_-GTP, 10 HEPES was used (Osmolarity: 283-289 mOsm, and pH: 7.2-7.4). The calculated liquid junction potential was –12 mV which is not corrected for in reported membrane potentials.

Whole-cell patch-clamp recordings were obtained using a Multiclamp 700B amplifier (Molecular Devices), digitized at 20 kHz (Digidata 1440, Molecular Devices) and recorded on a PC using pClamp 10 software (Molecular Devices). Input capacitance (C_m_) was measured using the Membrane Test protocol implemented in pClamp. Specifically, the electrode capacitance was compensated following gigaseal formation prior to breaking the membrane for establishing the whole-cell configuration. Under whole-cell voltage clamp mode (holding potential at −68 mV), a continuous square wave voltage command (5 mV, 10 ms) was applied at 50 Hz. Signals were low-pass filtered at 10 kHz and averaged 100 times. Recordings are obtained from cells with access resistance below 20 MΩ. The average access resistance were similar between control and 21RRS (Control vs. 21RRS: 8.67 ± 0.43 MΩ n = 72 vs. 7.58 ± 0.38 MΩ n = 67, p > 0.05, unpaired t-test). Resting membrane potential was measured in I = 0 current clamp mode around 2 min after breaking the membrane. ACSF contained kynurenic acid (2 mM, Sigma) and picrotoxin (100 uM, Sigma) to block ionotropic glutamate and GABA receptors, respectively except for glutamate- and synaptically-evoked excitation experiments specified below.

In order to examine neural excitation in response to an excitatory neurotransmitter glutamate, L-glutamic acid (5 mM, Sigma) was pressure applied using a Picospritzer II (10 psi, 500 ms, Parker, Pine Brook, NJ) from a glass pipette, with the tip placed 20-50 μm away from the soma of the recorded neurons. For individual trials, the location of the pipette tip was adjusted such that a single application generated varying amplitudes (∼50, 100 and 200 pA) of peak current response in the voltage-clamp. Thereafter, neural excitation was examined under the current clamp. The membrane potential prior to glutamate application was set near −55 mV, just below action potential threshold. Picrotoxin (100 uM, Sigma) was added to aCSF for these experiments.

In order to examine postsynaptic spike responses to synaptically-driven excitation, the afferent synapses were stimulated at various frequencies (5, 10 and 20 Hz for 2 sec) using an extracellular electrode placed 50-100 μm away from the postsynaptic neurons. For individual trials, we first recorded excitatory postsynaptic current (EPSCs) in voltage clamp configuration. We then performed current clamp experiments to examine excitatory postsynaptic potential (EPSP)-spike coupling by delivering the trains of synaptic stimulation. Prior to the synaptic stimulation, the membrane potential was held near −55mV. Picrotoxin (100 uM) was added to aCSF for these experiments.

For offline analysis of electrophysiological data, Clampfit 10 software (Molecular Devices, San Jose, CA) was used to measure the delay to first spike and frequency of firing. Resting membrane potential was determined from the peak of the histogram of the data points from an epoch of 10 second gap-free (I=0 mode) recording. The spike amplitude, threshold, rise time, half-width, and after-hyperpolarization peak were calculated using MiniAnalysis software (Synaptosoft, Fort Lee, NJ) for the first spike at the rheobase current injection.

### Electron microscopy

Mice were anesthetized with a mixture of pentobarbital (100 mg/kg, intraperitoneally) and transcardially perfused with PBS followed by 3.5% acrolein and 4% PFA. Fifty-micrometer-thick coronal sections of the brains were cut using a vibratome (Leica Biosystems) and stored at −20°C in cryoprotectant until further processing.

Brain sections were rinsed in PBS (50 mM, pH 7.4) and then quenched with 0.3% hydrogen peroxide (H_2_O_2_) for 5 min followed by 0.1% sodium borohydride (NaBH_4_) for 30 min. Afterwards, sections were rinsed three times in PBS and incubated for 1h in blocking buffer (10% goat serum, 3% bovine serum albumin, 0.01% Triton X-100) and then overnight with a primary anti-RFP antibody (1:10000 dilution, Rockland). The next day, sections were rinsed three times in TBS and incubated for 2h with secondary antibody conjugated to biotin (1:300, Jackson ImmunoResearch Laboratories) and then for 1h with Vectastain® Avidin-Biotin Complex Staining kit (Vector Laboratories). Sections were developed in a Tris buffer solution (TBS; 0.05M, pH 7.4) containing 0.05% diaminobenzidine and 0.015% H_2_O_2_ and then rinsed with PBS.

Brain sections containing the PVN from Bregma −0.58 mm to −0.94 mm were used and processed for electron microscopy as described elsewhere (Tremblay et al., 2010). Briefly, first, brain sections were incubated for 1h at room temperature in 4% osmium tetroxide (EMS) combined to 3% potassium ferrocyanide in 0.1M PB. After washing with ddH2O, brain sections were incubated in a 1% thiocarbohydrazide solution (in ddH2O) for 20 min at room temperature. A second incubation in 2% osmium tetroxide (in ddH2O) was next performed for 30 min at room temperature. After washing, sections were dehydrated in increasing concentrations of ethanol, treated with 100% propylene oxide, and impregnated in Durcupan ACM resin (EMS). Finally, the tissue sections were mounted between ACLAR sheets (EMS), embedded in a thin layer of resin, and placed in a 55°C oven for 3 days of polymerization.

The areas of interest containing the PVN were cut at 70 nm with an ultramicrotome (Leica Ultracut UC7, Leica Biosystems). Ultrathin sections were collected on square-mesh grids and examined at 80kV using a FEI Tecnai Spirit G2 microscope. For analysis, PVN-CRH neurons were randomly photographed in each mouse at the sequential magnifications of 8900x, 1900x, and 4800x using an ORCA-HR camera (10 MP; Hamamatsu).

Membrane contours of PVN-CRH neurons (n=7-10 neurons in each of 4 control mice and 4 restraint stressed mice) were drawn using the freehand selections tool of ImageJ 1.6 software. The area and perimeter, as well as shape descriptors “circularity”, “aspect ratio”, “roundness” and “solidity” were determined for each neuron.

### Multiphoton Imaging

Multiphoton imaging during electrophysiology recordings was performed with a multiphoton microscope (Bergamo II, Thorlabs Imaging Systems, Sterling, VA) using a 20× objective lens (XLUMPLFLN, 1.0 NA, water immersion, Olympus, Tokyo, Japan). Recorded cells were visualized by including Alexa Fluor 488 hydrazide (0.2 mM, ThermoFisher Scientific) in the internal solution. The images were acquired with the following settings: 16-bit dynamic range, 2× cumulative image averaging (3.8 fps), 0.32 × 0.32 µm pixel size, 0.7 μm z-axis interval. 750 nm laser (Chameleon, Coherent Laser Group, Santa Clara, CA) was applied, and the emission signals were detected through 562 nm edge epi-fluorescence dichroic mirror (FF562-Di03-32×44-FX, Semrock, Rochester, NY) and 525/50 nm single-band bandpass filter (FF03-525/50-25, Semrock). Each z-stack was obtained with 30 sec intervals. The surface membrane area was measured using Imaris software as described for the analysis for the biocytin-filled cells.

### Experimental design and Statistical analysis

All statistical analysis were performed using the GraphPad Prism 7 (Graphpad Software Inc., San Diego, CA). For c-Fos immunohistochemistry, two to four images of the PVN were obtained per animal, and the average value for an individual animal was considered as a N of 1. For electrophysiology, reconstruction of biocytin-filled neurons, electronmicroscopy and multiphoton imaging data, an individual cell/recording was considered as a n of 1 for the statistical analysis. The number of animal in a treatment group as shown as N. To perform a two-group comparison, an unpaired t-test and Mann-Whitney test were used for data sets for which a Gaussian distribution was assumed and not, respectively. To compare multiple groups, a one-way ANOVA was performed. Two-way ANOVA was used to conduct multiple-factor comparisons. To examine linear correlations, a linear regression analysis was conducted. p < 0.05was considered statistically significant.

## Results

### PVN-CRH neurons habituate to repeated stress

Exposure to a single restraint stress strongly activates the HPA axis, but this stereotyped neuroendocrine response diminishes after repeated exposures to the same stressor (Kiss and Aguilera, 1993; Bhatnagar et al., 2002). This indicates that the HPA axis habituates to repeated restraint (Grissom and Bhatnagar, 2009) and offers a model to study neural plasticity mechanisms that contribute to the decrease of stress responsiveness. We first characterized the time course of neuronal habituation in mice by studying the loss of PVN-CRH neurons response to repeated restraint. Using immunohistochemistry, we examined the expression of c-Fos, a marker for recent neuronal activity (Hoffman et al., 1993; Kovács, 1998; Girotti et al., 2006), in a CRH reporter line that was validated for specific fluorescent reporter (tdTomato) expression in CRH neurons in the PVN (Wamsteeker Cusulin et al., 2013; Chen et al., 2015). C-Fos expression was examined immediately after a 1 hour-long episode of restraint stress that was given for the first time (Acute), one day after 7 or 21 (1 hour-long) episodes of daily repeated restraint (7RRS or 21RRS), or one day after 7 days of a no stress recovery period following 21RRS (Recovery). As a control, we also studied stress-naïve (Control) mice (**Figure 1A**). As expected, acute restraint strongly increased c-Fos expression in PVN-CRH neurons as compared to control (% area for c-Fos/tdTomato; F (4, 27) = 44.5, p = < 0.0001, one-way ANOVA; Tukey’s post-hoc test for Control vs. Acute: 1.84 ± 0.69%, N = 5 vs. 38.21± 2.45%, N = 8; p < 0.0001; **Figure 1B, C**). 7RRS was not sufficient to decrease the restraint-induced c-Fos expression (7RRS: 35.44± 2.83%, N = 6; p = 1.0, vs. Acute). By contrast, 21RRS significantly decreased the c-Fos induction (21RRS: 11.91± 1.94%, N = 8; p < 0.0001, vs. Acute) to a level not statistically different from no-stress control (p = 0.066, 21RRS vs. Control). The loss of restraint-induced c-Fos by 21RRS was reversed after the 7 days of no-stress period to a level similar to acute restraint (Recovery: 36.90 ± 3.06%, N = 6; Recovery vs. Acute, p = 1.0). These results demonstrate that 21RRS, but not 7RRS, decreases the responsiveness of PVN-CRH neurons to the repeated stressor in a reversible manner. We then measured plasma corticosterone levels in the same groups of mice and found a significant positive correlation between the magnitude of c-Fos expression in PVN-CRH neurons and circulating corticosterone levels (R^2^ = 0.37, p < 0.0001, **Figure 1 D**). These results establish the loss of stress responsiveness in PVN-CRH neurons as a neural correlate for HPA axis habituation to repeated restraint.

### Repeated restraint stress decreases the intrinsic excitability of PVN-CRH neurons

Because intrinsic excitability directly influences firing activity, we reasoned that one simple explanation for the loss of c-Fos induction in PVN-CRH neurons after 21RRS would be a decrease in the intrinsic excitability of these neurons. To test this, we conducted whole-cell patch clamp recordings from PVN-CRH neurons in brain slices prepared one day after 21RRS without subjecting the mice to another restraint on the day of sacrifice (i.e. 24 h after the 21^st^ restraint). This time point was chosen to study the excitability of PVN-CRH neurons *ex vivo*, with relevance to their responsiveness to a stimulus that excites them *in vivo* (i.e. restraint). As a control, we also recorded from age-matched, stress-naïve mice. In the current-clamp configuration, PVN-CRH neurons were depolarized to elicit action potential firing with incremental current injection steps (10 pA for 700 ms) following a brief hyperpolarizing pre-pulse (−20 pA for 300 ms, **Figure 2A**). We found that 21RRS caused a strong leftward shift in the input-output (i.e. current injection-firing frequency) relationship (**Figure 2B, C**), revealing a robust decrease in intrinsic excitability. If this decrease in intrinsic excitability contributed to the *in vivo* changes in the PVN-CRH neurons’ responsiveness to the last restraint (as measured by c-Fos induction), the time course of this neural plasticity should match that of the c-Fos response. Indeed, we found that neither a single acute stress nor 7RRS were sufficient to change the input-output relationship (**Figure 2C**). Moreover, the 7-day no stress recovery period following 21RRS resulted in a spontaneous recovery of the excitability (**Figure 2C**). **Figure 2D** re-plots the firing frequency at one of the current injection steps (50 pA) shown in **Figure 2C**, clarifying that only 21RRS significantly decreased the firing frequency (F (4, 271) = 10.18, p < 0.0001, one-way ANOVA; Tukey’s post-hoc test for Control vs. 21RRS: 29.11± 5.23 Hz, n=72 vs. 23.03 ± 5.65 Hz, n = 67; p < 0.0001). These results show that repetitive firing frequency decreases with a time course similar to the habituation of c-Fos response *in vivo*.

**Figure 2.**
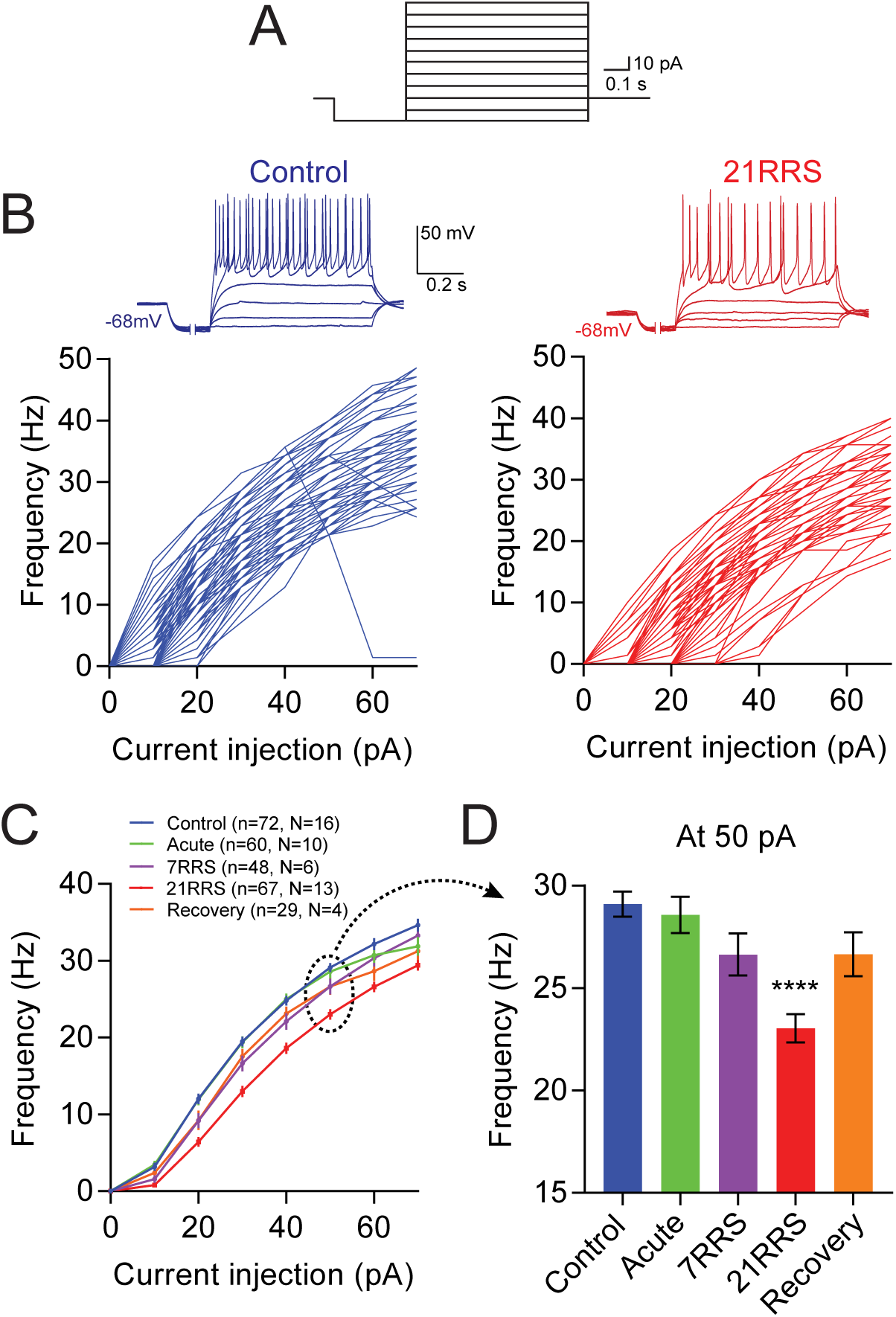
Firing frequency of PVN-CRH neurons decreases after 21RRS, but not 7RRS. **(A)** Current injection protocol (10 pA increment from ‒20 pA, holding potential = ‒68 mV). **(B)** Top, sample traces recorded from PVN-CRH neurons in slices from control (blue) and 21RRS (red) mice. Note that traces during the initial hyperpolarization step (0.3 s) are shortened (indicated by two vertical lines) for presentation purpose. Bottom, firing frequency-current curve of individual cells recorded in slices from control and 21RRS mice. **(C)** Graphs of average firing frequency-current curve. **(D)** Summary bar graph of frequency at 50pA current injection derived from data shown in C. Only 21RRS group significantly decreased firing frequency as compared to Control group. **** p < 0.0001. Error bars are S.E.M.

A recent study reported that PVN-CRH neurons increased firing latency (the delay before the first spike) due to voltage-dependent rapidly-activating and rapidly-inactivating K^+^ currents (I_A_) after a single acute swim stress in mice (Senst et al., 2016). Similar to this, we found that 21RRS had a trend toward increasing firing latency compared to control (F (4, 271) = 2.398, p = 0.051, one-way ANOVA; Tukey’s post-hoc test for Control vs. 21RRS: 0.14± 0.0075 s vs. 0.17 ± 0.012 s; p = 0.17; **Figure 3A-C**). Importantly, our current clamp protocol favors the I_A_-mediated firing latency as it includes a hyperpolarizing pre-pulse (−20 pA for 300 ms) that removes the inactivation of I_A_ (Yellen, 2002; Cai et al., 2007) before the depolarizing current steps. However, the time course of the development of stress-induced firing latency did not follow that of the c-Fos response (**Figure 3C**). That is, firing latency did not substantially increase with daily repeated challenges (Acute vs. 7RRS: p = 1.0; Acute vs. 21RRS: p = 0.98). The lack of correlation between c-Fos response and firing latency suggests that the stress-induced increase in firing latency does not account for c-Fos response habituation to 21RRS, and indirectly supports the importance of the decrease of repetitive firing frequency as a neuronal correlate for HPA axis habituation.

**Figure 3.**
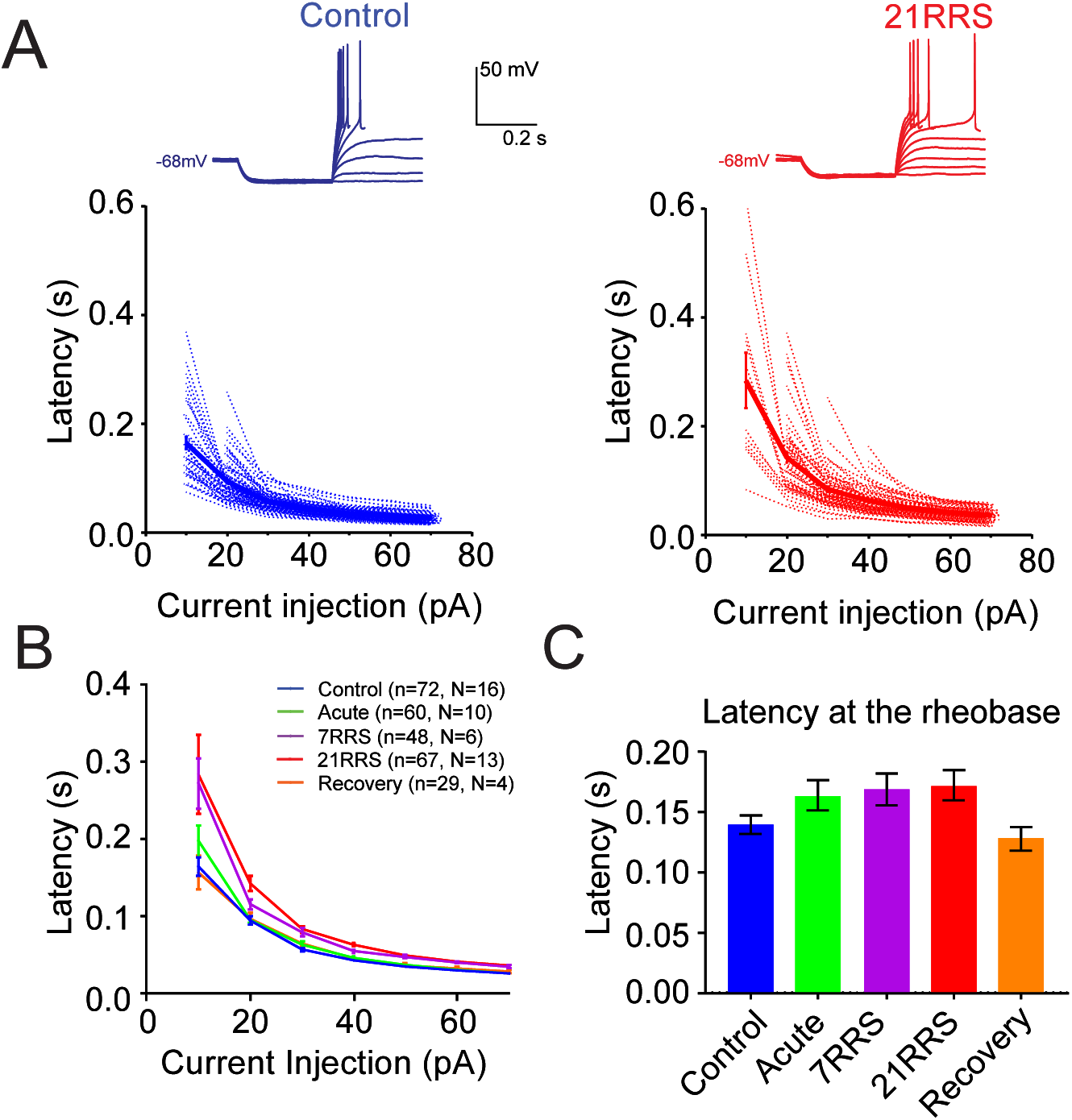
Firing latency of PVN-CRH neurons increases with an acute stress and does not further increase with repeated stress. **(A)** Top: sample traces depicting firing latency recorded from PVN-CRH neurons in slices from control (blue) and 21 day repeatedly stressed (red) mice. Bottom: firing latency-current curves of individual cells. **(B)** Graphs for average firing latency-current curves. **(C)** Summary bar graph of firing latency (at the rheobase current). Error bars are S.E.M.

### Repeated restraint stress decreases the responsiveness of PVN-CRH neurons to synaptic excitation

Glutamate synapses are the major afferent excitatory inputs to PVN-CRH neurons (van den Pol et al., 1990) and mediate the HPA axis response to restraint stress *in vivo* (Ziegler and Herman, 2000). To test whether 21RRS decreases the firing of PVN-CRH neurons in response to physiologically relevant stimuli, we examined the responsiveness of PVN-CRH neurons to focal application of glutamate (5 mM L-glutamic acid). In voltage-clamp configuration, we first set the location of the glutamate application glass pipette to generate transient postsynaptic current responses with peak amplitudes near 50 pA and 100 pA in individual cells (**Figure 4A**). We found that the resulting charge transfer was similar between control and 21RRS groups (F (1, 40) = 0.67, p = 0.42 for stress, two-way ANOVA), validating that the focal glutamate application caused similar amount of excitatory current influx between control and 21RRS groups. We then recorded in current-clamp mode and examined the firing responses to glutamate application. We found that the number of action potentials elicited by the focal glutamate applications was significantly reduced in 21RRS compared with control groups (F (1, 40) = 14.89, p = 0.0004, F (1, 40) = 8.59, p = 0.0056 and F (1, 40) = 0.18, p = 0.67 for stress, peak current and interaction, respectively; two-way repeated ANOVA; **Figure 4C**). These results suggest that 21RRS decreases the firing activities of PVN-CRH neurons in response to glutamate-mediated excitation.

**Figure 4.**
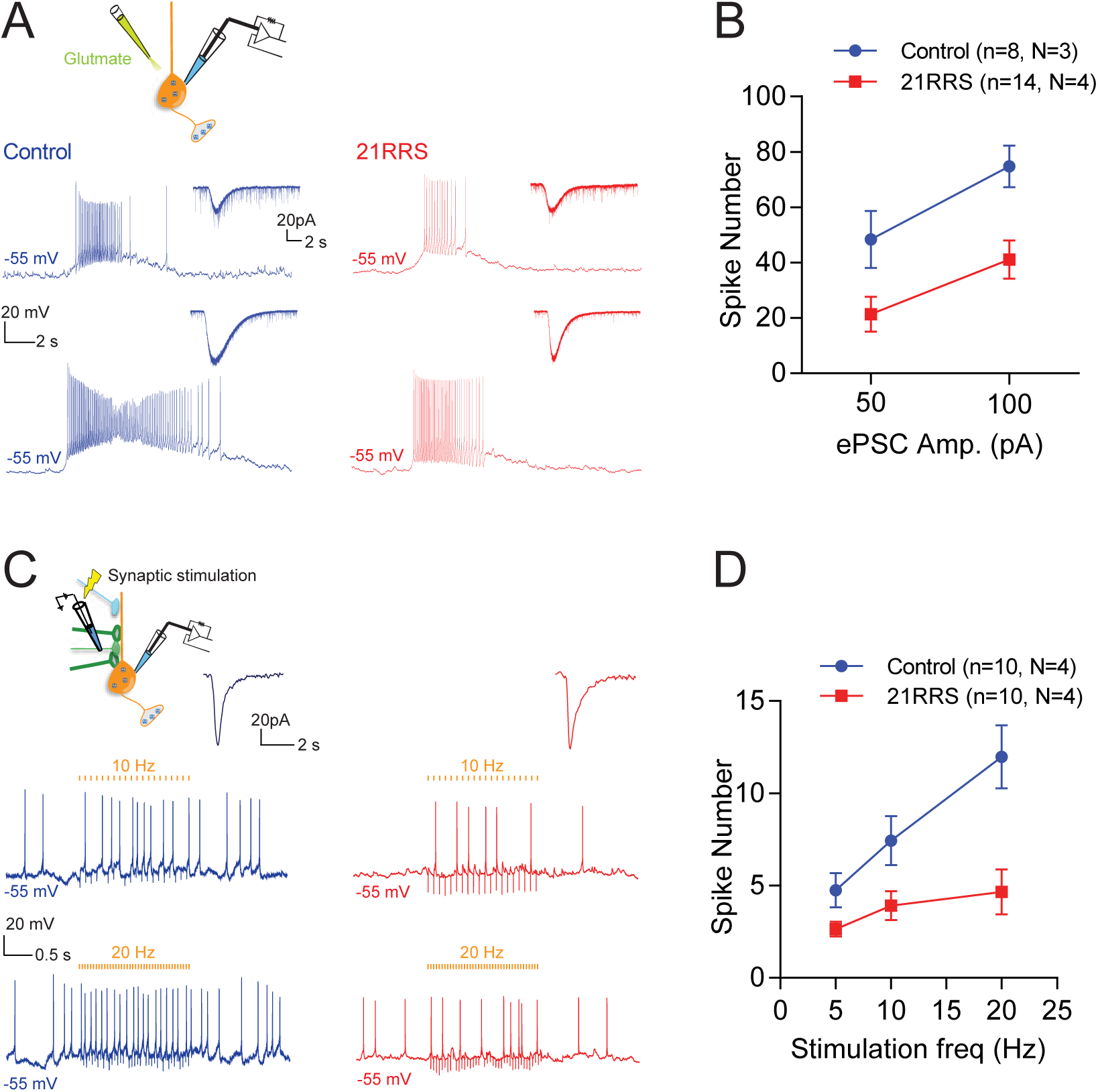
Repeated restraint stress diminishes firing of PVN-CRH neurons in response to glutamatergic excitation. **(A)** Top: a schematic for focal glutamate application. Bottom: focal glutamate application was adjusted to generate 50 pA and 100 pA inward current amplitude in voltage clamp (inset traces). In the same cells, firing response was examined in current clamp (bottom traces). **(B)** Summary of cell firing in response to focal glutamate applications (counted during the first 1 s after the onset of glutamate application). **(C)** Top: a schematic for synaptic stimulation. Bottom: postsynaptic currents after synaptic stimulation were first recorded (inset traces). In the same cells, spike responsse to different frequencies of synaptic stimulation were recorded (bottom traces). **(D)** Summary graph of spike number in response to increasing frequencies of synaptic stimulation. Error bars are S.E.M.

To address whether the decreased intrinsic excitability affects synaptically-induced firing of PVN-CRH neurons, we examined excitatory postsynaptic potential (EPSP)-spike coupling. We stimulated afferent inputs at 5, 10 and 20 Hz using an extracellular glass electrode in the presence a GABA_A_ receptor antagonist picrotoxin (100 μM) in order to pharmacologically isolate fast glutamatergic synaptic transmission. 21RRS significantly reduced EPSP-spike coupling across all stimulation frequencies (F (2, 36) = 20.98, p < 0.0001, F (1, 18) = 9.601, p = 0.006, and F (2, 36) = 7.113, p = 0.003 for stimulation frequency, stress and interaction, respectively; two-way repeated ANOVA; **Figure 4C, D**). In the same set of neurons, we found no difference in the amplitude of evoked excitatory postsynaptic currents (eEPSCs) and paired-pulse-ratio (PPR) recorded in voltage-clamp mode between control and 21RRS groups (data not shown). These results revealed that 21RRS significantly decreased firing response to synaptically-delivered excitation in PVN-CRH neurons.

### Repeated restraint stress decreases input resistance

We next investigated the cellular mechanisms underlying the stress-induced decrease of intrinsic excitability. When analyzing the current clamp data shown in **Figure 2**, we noticed that 21RRS strongly diminished the sub-threshold voltage responses to current injection steps (**Figure 5A**), pointing to changes in the passive membrane properties. Indeed, we found that the slopes of the voltage-current relationship were near linear in the subthreshold range for both groups and were less steep in 21RRS than control (**Figure 5B**). Consequently, the input resistance, calculated from the slope between −10 pA and 0 pA was significantly lower after 21RRS than control (Control vs. 21RRS: 1.30 ± 0.052 GΩ, n=72, vs. 1.04 ± 0.042 GΩ, n=67; p = 0.0001, unpaired t-test, **Figure 5C**). In line with this change, 21RRS significantly increased the rheobase (the amount of current injection required to trigger the first spike) (Control vs. 21RRS: 15.28 ± 0.66 pA, n=72, vs. 21.34 ± 0.97 pA, n=67; p < 0.0001, Mann-Whitney test, **Figure 5D**). On the other hand, 21RRS did not change the spike amplitude, threshold, rise time, half-width and after-hyperpolarization peak (AHP) (**Table 1**). We also found that the proportion of neurons that fired action potentials at the resting membrane potential was significantly lower in 21RRS than control (Control vs. 21RRS: 70.83% vs. 50.75%, p = 0.015, Chi-square test, **Figure 5E**). However, the resting membrane potential of neurons that did not fire was similar between control and 21RRS groups (Control vs. 21RRS: −55.76 ± 2.01 mV, n = 21, vs. −56.65 ± 1.08 mV, n = 34; p = 0.67, unpaired t-test; Table 1). These data collectively indicate that membrane properties that altered the input resistance contributed to the repeated stress-induced decrease of repetitive firing.

**Figure 5.**
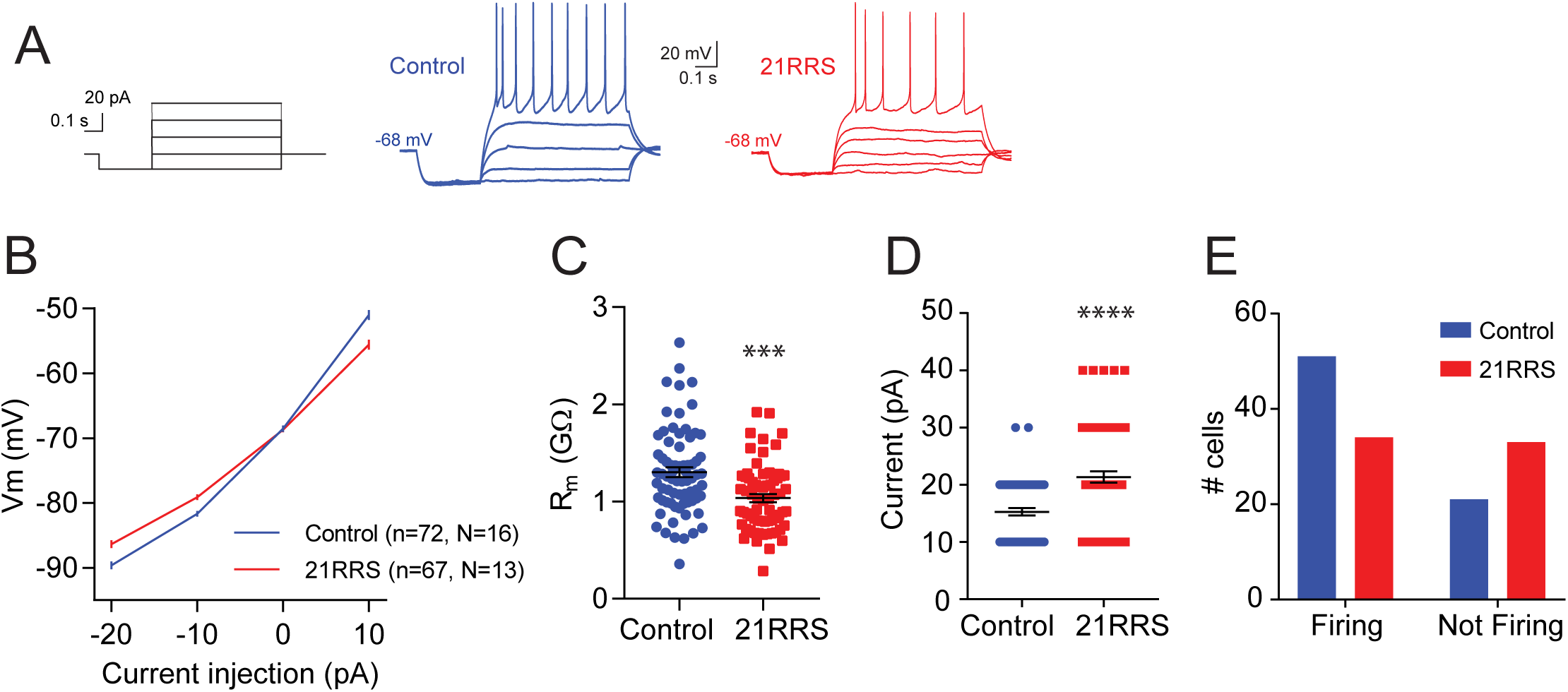
Repeated restraint stress decreases input resistance, increases rheobase and decreases proportion of cells firing at resting membrane potential (RMP). **(A)** Current injection protocol (10 pA increment from ‒20 pA, black), sample traces from control (blue) and 21RRS (red). **(B)** Summary graphs of the voltage-current relationship. **(C)** A summary graph of the input resistance calculated from the slope between ‒10 pA and 0 pA shown in B. *** p = 0.0001 **(D)** A summary graph of rheobase. **** p < 0.0001 **(E)** A comparison of the proportion of cells firing vs. not firing at resting membrane potential (RMP) for control and 21RRS groups; control cells fire more at RMP compared to 21RRS. Error bars are S.E.M.

**Table 1.**

| Properties | Control<br>(n = 72, N = 16) | 21RRS<br>(n = 67, N = 13) | p values |
| --- | --- | --- | --- |
| Action potential amplitude (mV) | 64.48 ± 0.79 | 66.11 ± 0.64 | 0.11 |
| Action potential threshold (mV) | -28.29 ± 0.48 | -27.98 ± 0.51 | 0.66 |
| Action potential rise time (ms) | 0.33 ± 0.0084 | 0.34 ± 0.0098 | 0.19 |
| Action potential half-width (ms) | 1.08 ± 0.025 | 1.14 ± 0.032 | 0.10 |
| After hyperpolarization (mV) | -19.89 ± 0.59 | -19.04 ± 0.54 | 0.29 |

### Voltage-dependent, non-inactivating input conductance correlates with firing frequency

To further investigate stress-induced changes in membrane properties that decreased the frequency of repetitive firing, subsets of PVN-CRH neurons shown in **Figure 2** were further studied in voltage-clamp mode. In this subset of neurons, we first used current-clamp recordings and confirmed that the firing frequency was significantly lower in 21RRS than control groups (Control vs. 21RRS: 29.73 ± 0.99 Hz, n = 32, vs. 23.57 ± 0.89 Hz, n = 38; p < 0.0001, unpaired t-test) which was accompanied by lower input resistance (Control vs. 21RRS: 1.49 ± 0.10 GΩ, n = 32, vs. 1.07 ± 0.06 GΩ, n = 38; p = 0.0003, unpaired t-test). Considering the prominent change in the input resistance, we hypothesized that 21RRS should affect sustained, non-inactivating input (whole-cell) currents when studied in voltage-clamp. Indeed, we found that the outward input currents measured at the end of incremental voltage steps (10 mV steps from −80 mV, for 10 s) were substantially larger in 21RRS than the control group (**Figure 6A, B**). **Figure 6C** plots input conductance calculated from the current-voltage relationship and revealed highly significant effects of 21RRS on voltage-dependent changes in the input conductance (F (1, 59) = 53.68, p < 0.0001; F (11, 649) = 751.7, p < 0.0001; and F (11, 649) = 54.31, p < 0.0001 for stress, voltage and interaction; two-way repeated ANOVA). Two important changes were noted. First, in line with the stress-induced decrease in input resistance found in the current-clamp experiment (**Figure 5**), the input conductance in the subthreshold range (more negative than −20 mV) was nearly constant (i.e. little voltage-dependent change) and had a trend to be higher in 21RRS than control groups (Control vs. 21RRS: p > 0.05 for −70 to −20 mV, two-way repeated ANOVA, Sidak’s post-hoc test): these changes did not reach statistical significance likely due to much greater changes at more positive membrane potentials (see below). Second, at potentials more positive than –10 mV, voltage-activated conductance became evident both in Control and 21RRS groups, revealing robust stress-induced differences (Control vs. 21RRS: p < 0.002 for −10 to + 30 mV, two-way repeated ANOVA, Sidak’s post-hoc test). We then plotted the correlation between firing frequency and input conductance calculated at individual voltage steps (from −70 to +30 mV) and found significant negative correlations at voltages more positive than 0 mV, with corresponding voltage-dependent increases in R^2^ values (**Figure 6D inset**). **Figure 6D** shows examples of the correlation at the largest (+30 mV) depolarization step (R^2^ = 0.45, p < 0.0001). These data show that 21RRS increased high-voltage activated, non-inactivating outward conductance, pointing to a potential mechanism that decreased repetitive firing frequency.

**Figure 6.**
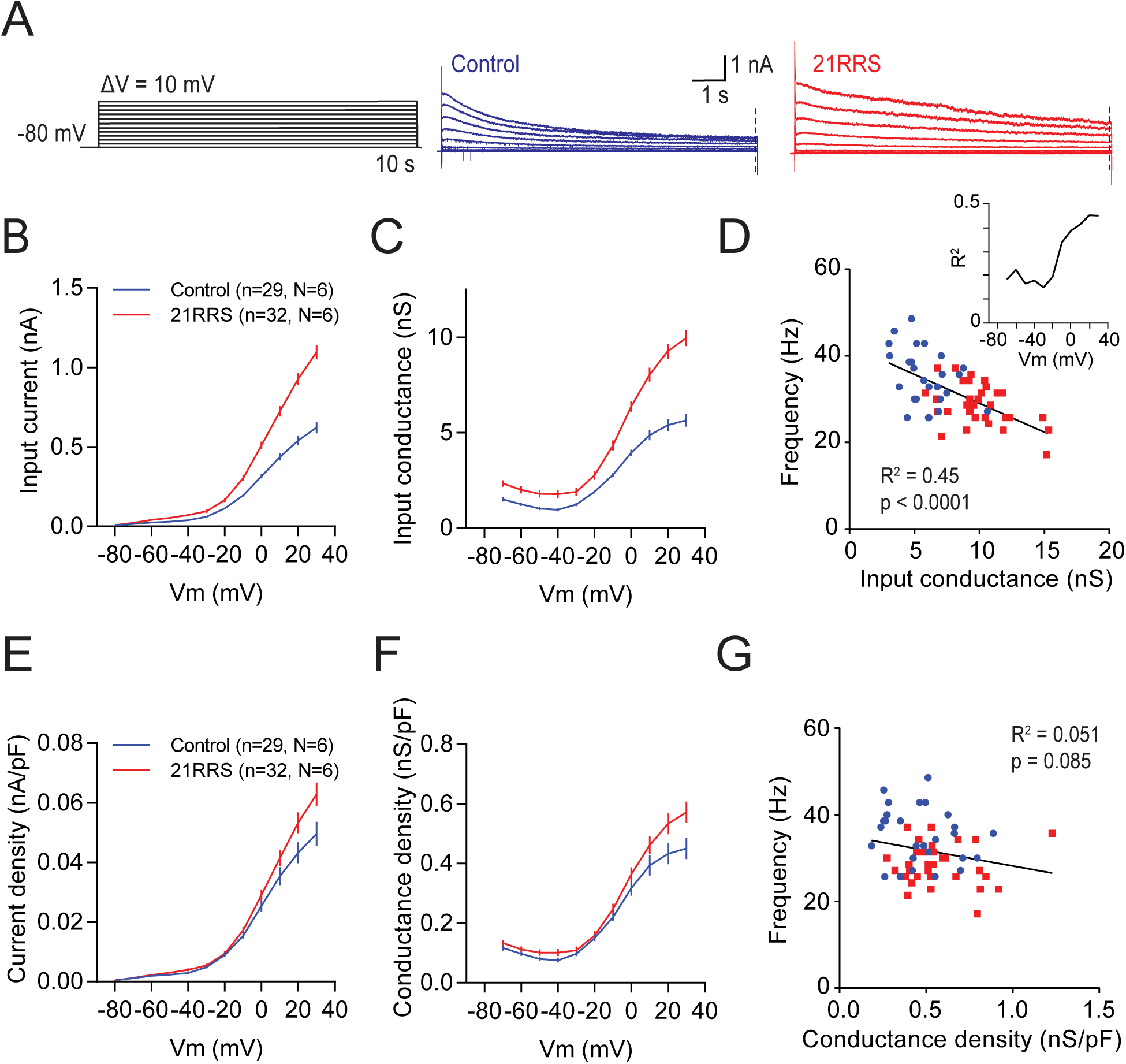
Input conductance, but not conductance density, correlates with firing frequency. **(A)** Voltage step protocol (+10 mV increment from −80 mV, black), sample traces for current responses from control (blue) and 21RRS (red) cell. Orange dotted line on sample traces indicates where non-inactivating input conductance was measured (t = 9.996 s after the onset of voltage steps). **(B)** Average input current-voltage curves for control and 21RRS. **(C)** Average input conductance-voltage curves. **(D)** Input conductance (measured at 30 mV) significantly correlated with the firing frequency. Inset. Summary of R^2^ values across different voltages. **(E)** Average current density-voltage curves for control and 21RRS. **(F)** Average conductance density-voltage curves. **(G)** Conductance density (measured at 30 mV) does not correlate with firing frequency. Error bars are S.E.M.

In addition to the sustained, non-inactivating conductance, 21RRS significantly increased rapidly inactivating outward conductance (measured 3 ms after the onset of voltage steps) that was activated upon depolarization (> −30 mV), an observation consistent with the stress-induced increase of I_A_. However, these changes did not correlate with firing frequency (R^2^ = 0.053, p = 0.076 vs. current at 30 mV). Similarly, slowly inactivating conductance (measured at 116 ms after the onset of voltage steps) was not altered by 21RRS (R^2^ = 0.020, p = 0.28 vs. current at 30 mV). Taken together, our data suggest that voltage-dependent, non-inactivating conductance played a major role in the stress-induced decrease of repetitive firing frequency.

### Repeated stress did not change conductance density

Input conductance is a product of the conductance density (conductance per unit membrane area) multiplied by the total surface membrane area (i.e. cell size). To evaluate the effects of 21RRS on the conductance density, we normalized the input current and conductance by cell capacitance (C_m_), an electrophysiological measure to estimate surface membrane area (Golowasch et al., 2009; Taylor, 2012), in individual cells (**Figure 6E, F**). Strikingly, the normalization largely eliminated the stress-induced changes (F (11, 649) = 367.7, p < 0.0001; F (1, 59) = 3.421, p = 0.069; and F (11, 649) = 4.437, p < 0.0001 for voltage, stress and interaction, respectively; two-way repeated ANOVA) and abolished the correlation between the firing frequency and the high-voltage activated, sustained conductance (R^2^ = 0.051, p = 0.085, **Figure 6G**). These results led us to examine potential changes in the size of PVN-CRH neurons and their contributions to intrinsic excitability.

### Repeated stress increased cell size

As predicted from the lack of difference in the conductance density between Control and 21RRS groups, C_m_ was significantly larger in 21RRS compared to controls (Control vs. 21RRS: 14.3 ± 0.45 pF, n = 73, vs. 19.0 ± 0.59 pF, n = 67; unpaired t-test; p < 0.0001, **Figure 7A**). Importantly, we found a significant negative correlation between C_m_ and firing frequency (R^2^ = 0.26, p < 0.0001, **Figure 7B**). This data suggested a scenario that PVN-CRH neurons grew their surface membrane area after repeated stress exposure. Indeed, in a rat model of chronic variable stress, a histological study has reported an increase in the soma size of neuroendocrine PVN-CRH neurons (Flak et al., 2009). Thus, we asked if 21RRS increases cell soma size of PVN-CRH neurons, and if so, whether such structural plasticity accounts for the observed increase in C_m_. To test this, we repeated another set of patch-clamp recordings followed by morphological reconstruction of the recorded neurons by including biocytin, a histology marker compatible with patch-clamp recording, in the internal solution (**Figure 7C**). First, we confirmed that 21RRS significantly decreased the firing frequency in this cohort (Control vs. 21RRS: 29.43 ± 1.16 Hz, n = 20, vs. 22.21 ± 1.14 Hz, n = 31; p < 0.0001, unpaired t-test), increased C_m_ (Control vs. 21RRS: 16.68 ± 0.97 pF, n = 20, vs. 20.95 ± 1.02 pF, n = 31; p = 0.0061; unpaired t-test), and that firing frequency and C_m_ significantly correlated (R^2^ = 0.27, p = 0.0001). In support of our prediction, histological measurements found that the cell soma area significantly increased after 21RRS (Control vs. 21RRS: 426.6 ± 16.29 µm^2^, n = 20, vs. 509.7 ± 15.91 µm^2^, n = 31; p = 0.001, unpaired t-test, **Figure 7C, D**). PVN-CRH neurons have relatively simple morphology mostly with two to three proximal dendrites (Rho and Swanson, 1989). While 21RRS did not change the number of primary dendrites (Control, 2.45 ± 0.17, n = 20, vs. 21RRS, 2.29 ± 0.13, n = 31; p = 0.50, Mann-Whitney test; **Figure 7E**), the diameter of primary dendrites increased significantly (Control vs. 21RRS: 26.23 ± 0.56 µm, n = 20, vs. 31.15 ± 0.79 µm, n = 31; p < 0.0001, unpaired t-test; **Figure 7F**), indicating that the stress-induced cellular hypertrophy is not limited to the cell soma. However, despite the robust hypertrophy of cell soma and proximal dendrites, the cell soma size did not significantly correlate with C_m_ (R^2^ = 0.0089, p = 0.50, **Figure 7G).**

**Figure 7.**
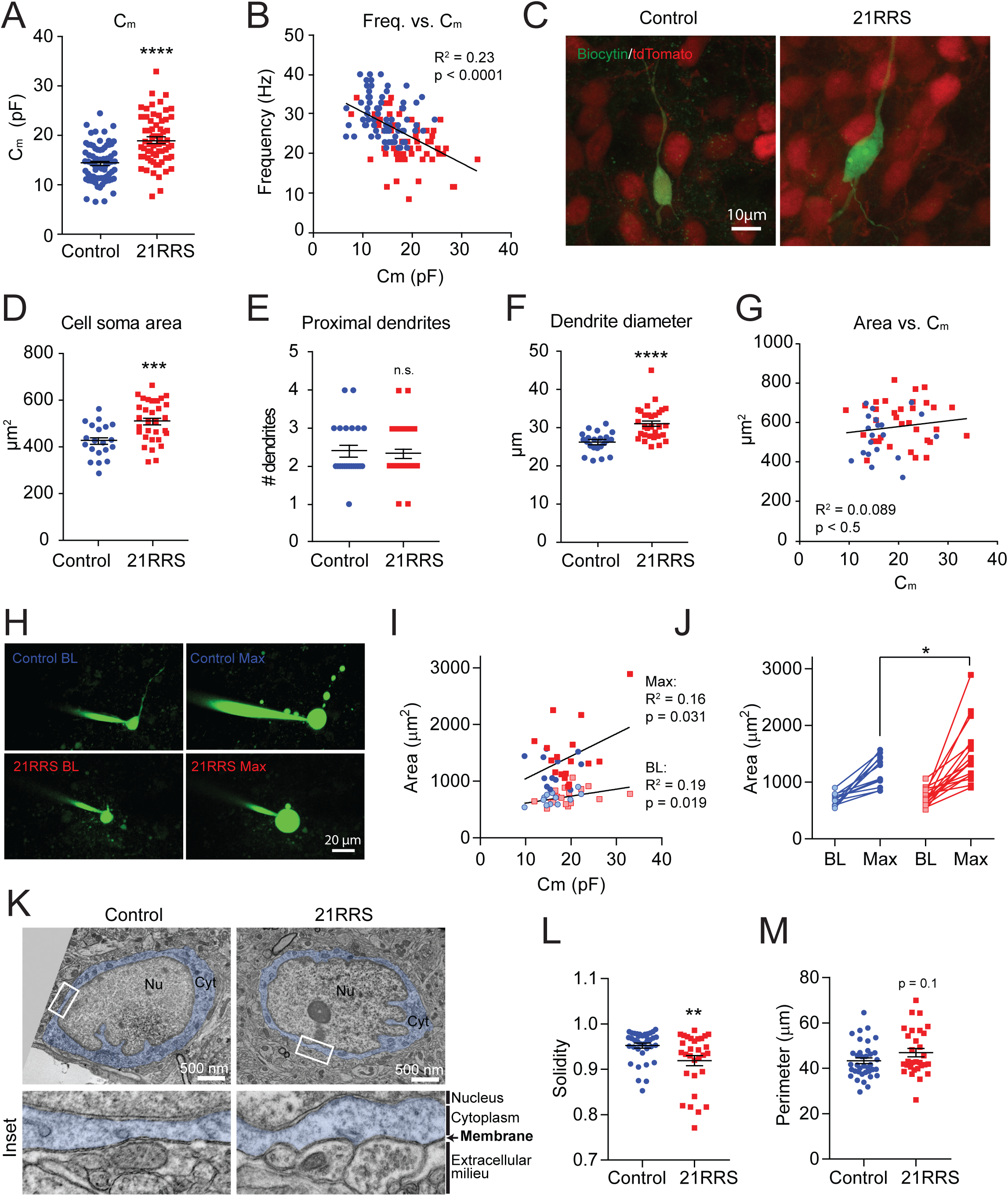
Repeated restraint stress increases C_m_ and causes hypertrophy of the cell soma and proximal dendrites. **(A)** Summary graph of C_m_ for control and 21RRS. **** p < 0.0001 **(B)** C_m_ negatively correlates with firing frequency. **(C)** Examples of biocytin-filled PVN-CRH neurons from control and 21RRS groups. **(D)** Summary graph of cell soma area measured from biocytin-filled PVN-CRH neurons. *** p = 0.001. **(E)** Summary graph of the number of the primary dendrites. **(F)** Summary graph of the diameter of primary dendrites (measured at 2 µm from the end of cell soma). **** p < 0.0001 **(G)** C_m_ does not correlate with cell soma area among biocytin-filled PVN-CRH neurons. **(H)** Examples of fluorescent dye-filled PVN-CRH neurons imaged using multiphoton microscopy. Examples of baseline (BL, left) and maximum swelling (Max, right) images for control (top) and 21RRS (bottom) cells. **(I)** Summary graph for the cell soma area before and after swelling by high osmolarity internal solution. Both control and 21RRS showed significant swelling (p < 0.0001, Two-way repeated ANOVA), and 21RRS cells swelled more than control (Control-Max vs. 21RRS-Max, * for p = 0.036, Sidak’s multiple comparison test). **(J)** Cm positively correlated with the soma surface area at baseline (R^2^ = 0.19, p = 0.019) and at maximum swelling (R^2^ = 0.16, p = 0.031). **(K)** Examples of immunohistochemical transmission electron microscopy of the soma of PVH-CRH neurons from control (left) and 21RRS (right) groups. Insets shows magnified images of the area indicated by white rectangles. **(L)** Summary graph for cell membrane perimeter of PVH-CRH neurons measured in transmission electron microscopy images (Unpaired t-test, p = 0.1). **(M)** Summary graph for solidity. ** p = 0.0092 (Mann-Whitney test). Error bars are S.E.M.

One possibility for the lack of correlation is the technical limitation associated with the reconstruction of cell soma after patch-clamp recording. That is, the retraction of the recording pipette from the soma after the electrophysiological recording inevitably removes variable amount of cell soma membrane. Thus, this introduces random deviation between the actual and reconstructed soma size. Another possibility is that the plasma membrane of mammalian cells is not smooth but has infoldings, ruffles and microvilli (Gentet et al., 2000). When measured using confocal imaging of biocytin-filled cells, these ultrastructural irregularities would cause an underestimation of the surface membrane area that contributes to C_m_ measurement. To address these limitations and further examine the relationship between cell soma size and C_m_, we imaged the morphology of fluorescent dye-filled PVN-CRH neurons using a multiphoton microscope during electrophysiological recordings (i.e. when the recording pipette was still attached to the cell soma). In addition, in an attempt to flatten out possible ruffles and infoldings of the membrane, we artificially expanded the plasma membrane of the recorded neurons and measured the maximum surface membrane area it can reach before rupture. Specifically, we took advantage of osmotic drive to expand neurons (Gentet et al., 2000) by obtaining a whole-cell patch clamp recordings with a high osmolarity (∼400 mOsm) internal solution. The “baseline” Z-stack image was measured immediately after the start of the whole cell configuration. The maximum size was measured by the last imaging before the cell rupture due to the osmotic pressure. In this subset of experiments, we confirmed that the 21RRS group had larger C_m_ than control group although this did not reach statistical significance likely due to relatively low sample number (Control vs. 21RRS: 16.32 ± 1.02 pF, n = 12 vs. 19.51 ± 1.17, n = 17, p = 0.061). Importantly, in this cell size measurement, we found a significant positive correlation between C_m_ and the soma surface area both at the baseline (R^2^ = 0.20, p = 0.016, **Figure 7I**) and at the maximum (R^2^ = 0.14, p = 0.043). These data indicate that the lack of correlation between C_m_ and cell soma area in the biocytin-filled neurons was due to the variability in the loss of membrane by pipet retraction as discussed earlier. Furthermore, we found that both control and 21RRS neurons significantly increased their soma size after the infusion of high osmolarity internal solution (**Figure 7H, J**). Importantly, this rapid swelling of neurons did not change C_m_ values that was measured in a pilot experiment (Baseline vs. Max, 20.51 ± 1.44 pF vs. 19.94 ± 1.24 pF, n = 12, p = 0.26, paired t-test). Thus, these data collectively indicate that PVN-CRH neurons have a substantial amount of plasma membrane ruffles/infoldings that biophysically contribute to the surface area (i.e. contribute to C_m_ and input resistance). Interestingly, cells from the 21RRS group were able to swell significantly more than the control group (F(1, 27) = 3.67, p = 0.066, F(1, 27) = 55.57, p < 0.0001, F(1, 27) = 2.30, p = 0.14 for stress, swelling and interaction; Sidak’s multiple comparisons test for Control-Max vs. 21RRS-Max, p = 0.037; **Figure 7J**). These data raised the possibility that, in addition to the gross cell soma size increase, the increase of surface membrane area after 21RRS is accompanied by an increase in membrane infoldings/ruffles that would increase the total membrane surface.

To measure potential changes in cell membrane ultrastructure at high spatial resolution, *in situ* in the intact hypothalamus, we used immunohistochemical transmission electron microscopy and obtained images of tdTomato-immunopositive neurons in the PVN (i.e. CRH-PVN neurons) of control and 21RRS groups. In line with our prediction, we found that the plasma membrane delineating the soma of PVN-CRH neurons became more ruffled with frequent small curvatures after 21RRS versus control (**Figure 7K**). This observation was supported by a significant decrease in the solidity that describes the extent of concavity in the structure after 21RRS (Control vs. 21RRS: 0.95 ± 0.0055, n = 37, vs. 0.92 ± 0.061, n = 30; p = 0.0092, Mann-Whitney test; **Figure 7L**). Other measures of changes in surface shape, including circularity, roundness and aspect ratio, did not reach statistical significance but showed trends that were consistent with an increase in concavity of the soma after 21RRS (**Table 2**). In line with the increase of cell soma size detected after 21RRS, we found that the perimeter of the soma showed a trend toward stress-induced increase although this change did not reach statistical significance (Control vs. 21RRS: 43.32 ± 1.27, n = 37, vs. 46.91 ± 1.84, n = 30; p = 0.1, unpaired t-test; **Figure 7M**). In summary, these data indicate that 21RRS increases ultrastructural irregularities in the plasma membrane that would increase the total membrane surface.

**Table 2.**

| Properties | Control<br>(n = 37, N = 4) | 21RRS<br>(n = 30, N = 4) | p values |
| --- | --- | --- | --- |
| Area (nm<sup>2</sup>) | 116.2 ± 5.34 | 122.5 ± 7.03 | 0.47 |
| Perimeter (nm) | 43.32 ± 1.27 | 46.91 ± 1.84 | 0.10 |
| Circularity | 0.78 ± 0.019 | 0.71 ± 0.025 | 0.07 |
| Aspect ratio | 1.53 ± 0.065 | 1.69 ± 0.080 | 0.12 |
| Roundness | 0.67 ± 0.026 | 0.63 ± 0.030 | 0.18 |
| Solidity | 0.95 ± 0.0055 | 0.92 ± 0.0011 | 0.0092 |

## Discussion

Habituation of the HPA axis response to recurrent stressors is a fundamental aspect of stress adaptation observed both in human and experimental animals (Aguilera, 1994; Kirschbaum et al., 1995; McEwen and Seeman, 1999; Uchida et al., 2008). The major finding of our study is that PVN-CRH neurons substantially decrease their intrinsic excitability over the course of daily repeated restraint, in a time course that coincides with their loss of stress responsiveness *in vivo*. Mechanistically, repeated stress increases cell size (i.e. surface membrane area): the direct consequence of this cell size increase is an increase in the input conductance that diminishes the cell’s voltage responses to the influx of positive charges (e.g. excitatory inputs). Moreover, our multi-photon and electron microscopy data reveal that the stress-induced cell size increase involves an augmentation of membrane ruffling/infolding; this ultrastructural plasticity may accommodate the surface membrane area increase with proportionally less expansion of gross cell volume. Taken together, we report a novel structure-function relationship for the plasticity of intrinsic excitability that correlates with the habituation of neuroendocrine response to repeated stress.

Previous studies showed that neuroendocrine habituation to repeated restraint was paralleled by a diminished c-Fos up-regulation in PVN-CRH neurons upon re-exposure to restraint (Melia et al., 1994; Stamp and Herbert, 1999; Girotti et al., 2006; Ons et al., 2010). Additionally, these studies also reported a similar reduction of c-Fos up-regulation in various stress-responsive brain areas including the amygdala, the prefrontal cortex and the lateral septum (Melia et al., 1994; Stamp and Herbert, 1999; Girotti et al., 2006; Ons et al., 2010). Thus, a prevailing idea is that the loss of PVN-CRH neurons’ response to repeated stress is primarily attributable to the diminished stress response in brain areas upstream of the PVN. Our study is the first to report, to the best of our knowledge, that the intrinsic excitability of PVN neurons decreases during neuroendocrine habituation to repeated stress. When challenged with a repeated stressor, the diminished activities of the brain areas upstream of the PVN would result in decreased excitatory synaptic inputs to (or excitation/inhibition balance at) the PVN-CRH neurons. Thus, we predict that the intrinsic plasticity of PVN-CRH neurons would work additively to synaptic changes, diminishing the overall responsiveness of the HPA axis.

We found a robust increase in the input conductance following repeated restraint. Voltage-clamp experiments revealed that the stress-induced increase of input conductance is most prominent at the end of long depolarizing voltage steps. Based on the voltage activation profile and the non-inactivating property, this membrane conductance underlies delayed rectifier (K_dr_) currents. K_dr_ have been reported in parvocellular neuroendocrine neurons in the PVN (Luther et al., 2002) and function to constrain the firing activity of neurons (Hoyda and Ferguson, 2010). Indeed, we found that firing frequency (measured in the current clamp) significantly correlated with this voltage-activated (> 20 mV), non-inactivating conductance characteristic of K_dr_ (Luther et al., 2002). In addition to K_dr_, we found that repeated stress increased I_A_, which is voltage-activated (> −40 mV) and rapidly inactivating (Yellen, 2002; Cai et al., 2007). However, this I_A_ did not correlate with the firing frequency, negating its major contribution to repetitive firing frequency. Senst et al. (2016) reported that a single swim stress rapidly increases firing latency through I_A_ in young (P22-35) mice (Senst et al., 2016). Our study, using restraint in adult mice (8-12 weeks), found a similar but less robust change that did not reach statistical significance. This difference in the magnitude could be due to differences in the age of mice and/or types of stressors used. Our study also extends this earlier observation by demonstrating that the stress-induced I_A_ persists but does not further develop during repeated restraint. A recent study reported another type of intrinsic plasticity in PVN-CRH neurons where chronic corticosterone treatment decreases repetitive firing frequency (Bittar et al., 2019). However, this hormone-induced plasticity did not involve changes in the input resistance or rheobase (unlike after repeated stress), pointing to the diversity the mechanisms that modulate the intrinsic excitability of PVN-CRH neurons in various stress-relevant conditions.

In addition to synaptic plasticity, ample data also supports the importance of experience-induced plasticity of intrinsic excitability in various neuronal types, species and learning paradigms (Daoudal and Debanne, 2003; Zhang and Linden, 2003; O’Leary et al., 2014; Kourrich et al., 2015). Common mechanisms for intrinsic plasticity are changes in conductance density (i.e. input conductance normalized by C_m_), attributable to corresponding up- and down-regulation of Na^+^, K^+^, and Ca^2+^ channels (Zhang et al., 1998, 2002; Dong et al., 2006; Kourrich and Thomas, 2009; Kourrich et al., 2015). By contrast, our study found little change in the conductance density (after normalization by C_m_). These results led us to investigate plasticity mechanisms that increased C_m_ and its relevance to the observed changes in the intrinsic excitability. In this study, we operationally defined C_m_ as the input capacitance, a product of the specific membrane capacitance (capacitance per unit area of membrane) multiplied by the total surface membrane area. Specific capacitance for plasma membrane (i.e. lipid bilayer) is commonly viewed as a biological constant near 1.0 µF/cm^2^ (Kandel et al., 2012) and has been empirically shown to be similar among different neuronal cell types (Gentet et al., 2000). While our data does not exclude the possibility that stress may change the specific conductance by changing lipid composition (Niles et al., 1988) and/or protein insertion into the plasma membrane (Valincius et al., 2008; Zimmermann et al., 2008), the simplest explanation for the observed increase in C_m_ is an increase in surface membrane area (Hodgkin and Huxley, 1952). We measured C_m_ in voltage-clamp by delivering a small subthreshold step. This method only measures the part of cell that is “well-clamped” and does not measure the total cell capacitance (Golowasch et al., 2009; Taylor, 2012). Thus, one limitation of our study is that the proportion of surface membrane area that contributes to the C_m_ measurement (i.e. “well-clamped”) remains unclear. We assumed that the soma and proximal dendrites that are electrotonically close to the point of voltage clamp would account for the major portion of C_m_. In line with this prediction, we found a significant stress-induced increase in the surface area of the soma in the morphological reconstruction of biocytin-filled neurons using confocal microscopy. Importantly, simultaneous C_m_ measurement by patch-clamp electrophysiology and cell soma surface area measurement using multiphoton microscopy found a significant positive correlation between the two measurements. Together, we conclude that cell soma size makes a major contribution to the change in C_m_. Herman and colleagues (2009) reported a similar increase in cell soma size of PVN-CRH neurons in a rat models of chronic variable stress (Flak et al., 2009). Our study revealed a biophysical consequence of such a stress-induced structural plasticity. In addition to the changes in cell soma size, we also found a significant increase in the diameter of the proximal dendrites, suggesting that dendritic hypertrophy also contributes to the observed change in C_m_. Interestingly, Francis et al. (2017) recently reported a complementary finding that chronic social defeat stress causes dendritic atrophy of dopamine D1 containing medium spiny neurons in the nucleus accumbens, and that this morphological plasticity was accompanied by a decrease in C_m_ and an increase in intrinsic excitability (Francis et al., 2017, 2019). These data raises an intriguing possibility that, in addition to ion channel modulations, changes in cell size (which is reflected by C_m_ and directly influences input conductance) may be a mechanism for experience-induced plasticity of intrinsic excitability generalizable beyond PVN-CRH neuron.

The actual surface membrane area of mammalian cells is often underestimated because the cell surface is not smooth due to various ultrastructural irregularities such as ruffles, infoldings and microvilli (Gentet et al., 2000; Zimmermann et al., 2008). A simple approach to smoothen out these surface irregularities is to swell the cell by manipulating the osmotic differences across the plasma membrane (Gentet et al., 2000; Zimmermann et al., 2008). By using hyperosmotic internal solution, our study showed that PVN-CRH neurons increased their cell soma surface area nearly 80% before they ruptured, indicating that PVN-CRH neurons have substantial surface irregularities that accommodate more surface membrane area than their gross cell size. Importantly, this relatively rapid swelling (ruptures around 5 min) did not increase C_m_, excluding an insertion of new membrane as occurs for slower adaptation to cell swelling (Dai et al., 1998; Zhang and Bourque, 2003). Interestingly, we also found that the magnitude of the swelling was greater in cells from 21RRS, suggesting that repeated stress increased surface irregularities. In line with this, our electron microscopy analysis found significantly more ultrastructural irregularities (ruffled membrane) in the cross sections of PVN-CRH neurons after 21RRS than control. While these increased surface irregularities might reflect a process of cell size expansion, it is tempting to speculate that they may also be a biological strategy to accommodate the surface membrane area increase with proportionally less expansion of gross cell volume. In summary, we report a novel structure-function relationship for the plasticity of intrinsic excitability that correlates with the habituation of the neuroendocrine response to repeated stress.

## Author contributions

SM designed experiments, acquired, analyzed and interpreted data, and drafted the article. AI, HI, XFW and ES acquired, analyzed and interpreted data. MH, NV, MET acquired, analyzed and interpreted electronmicroscopy data. WI conceived, designed experiments and drafted the article.

## Acknowledgements

We thank all members of Inoue lab for thoughtful inputs to the project. We are grateful to Ms. Irma Meteluch for her help with animal husbandry. We also thank Dr. Jaideep Bains (University of Calgary) for his constructive comments on early versions of the manuscript.

